# Fisheries genomics of snapper (*Chrysophrys auratus*) along the western Australian coast

**DOI:** 10.1101/2022.02.17.479830

**Authors:** Andrea Bertram, David Fairclough, Jonathan Sandoval-Castillo, Chris Brauer, Anthony Fowler, Maren Wellenreuther, Luciano B. Beheregaray

## Abstract

The efficacy of fisheries management strategies depends on stock assessment and management actions being carried out at appropriate spatial scales. This requires understanding of spatial and temporal population structure and connectivity, which is challenging in weakly structured and highly connected marine populations. We carried out a population genomics study of the heavily exploited snapper (*Chrysophrys auratus*) along ∼2,600 km of the Australian coastline, with a focus on Western Australia (WA). We used 10,903 filtered SNPs in 341 individuals from eight locations to characterise population structure and connectivity in snapper across WA and to assess if current spatial scales of stock assessment and management agree with evidence from population genomics. Our dataset also enabled us to investigate temporal stability in population structure as well as connectivity between WA and its nearest, eastern jurisdictional neighbor. As expected for a species influenced by the extensive ocean boundary current in the region, low genetic differentiation and high connectivity was uncovered across WA. However, we did detect strong isolation by distance and genetic discontinuities in the mid-west and south-east. The discontinuities correlate with boundaries between biogeographic regions, influenced by on-shelf oceanography, and the sites of important spawning aggregations. We also detected temporal instability in genetic structure at one of our sites, possibly due to interannual variability in recruitment in adjacent regions. Our results partly contrast with the current spatial management of snapper in WA, highlighting the need for a review. This study supports the value of population genomic surveys in informing the management of weakly-structured and wide-ranging marine fishery resources.

## 1 INTRODUCTION

Marine ecosystems are the last on the planet in which wild populations are heavily exploited for human consumption. The continual improvement of fisheries management practices is necessary to facilitate ongoing sustainable exploitation or recovery of overfished populations (Garcia and Ye 2018). The success of such efforts requires that stock status estimation and the quantitative prediction of the effects of different management choices are done accurately, and that applied management actions are effective (Hilborn and Walters 1992).

These matters depend upon the collection and evaluation of data and the implementation of regulations to fishing being done at appropriate spatial scales, requiring knowledge of stock structure and the levels of connectivity between stocks. Despite this, stock structure and connectivity are commonly ignored or not investigated. Instead, exploited fishes are often assessed and managed according to politically or geographically distinct areas (Reiss et al. 2009). The consequences of ignoring stock structure and connectivity in assessment and management have been widely documented (Fu and Fanning 2004, Sterner 2007, Kell et al. 2009, Ying et al. 2011, Kerr et al. 2014).

The biological characteristics of marine organisms that shape population structure and connectivity include habitat specialization determined by breeding site preferences, dietary requirements and abiotic tolerance limits (Cowen 2002, Cowen and Sponaugle 2009), and dispersal potential (Bohonak 1999) determined by reproductive strategy (e.g. broadcast spawning and brooding), the duration of any pelagic stages and the movement behaviour of both larval and post-larval stages. For example, in the broadcast spawning and highly migratory yellowfin tuna (*Thunnus albacares*), homogeneity likely occurs at spatial scales as extensive as entire ocean basins (Barth et al. 2017, Pecoraro et al. 2018). Due to the fluidity of the marine environment and processes like currents, waves and tides, connectivity and population homogeneity can occur over large spatial scales, particularly in species with long pelagic larval stages (Selkoe et al. 2008, Cowen and Sponaugle 2009). However, marine environments are dynamic and heterogeneous and therefore contain elements that disrupt connectivity and act as barriers to dispersal, including meanders and eddies, fronts, irregular coastline topology, habitat heterogeneity and countercurrents (Cowen 2002, Cowen and Sponaugle 2009). For example, Taillebois et al. (2017) detected significant population structuring in the broadcast spawning marine teleost, the black-spotted croaker (*Protonibea diacanthus*) across topographically complex coastline in northern Australia at spatial scales of only 100s of kilometres.

Our focus here is on snapper (*Chrysophrys auratus*), a large, long-lived demersal sparid distributed in the coastal waters of the southern two-thirds of Australia, as well as in northern New Zealand (Gomon et al. 2008). In Western Australia (WA), snapper supports highly valuable commercial and recreational fisheries from the Shark Bay region to the border of South Australia (SA; Figure 1), with the largest catches landed off the west coast (Gaughan and Santoro 2021). Across this region, snapper is used as an indicator species for the inshore suite of demersal scalefish resources. In WA, snapper is assessed and managed as three separate stocks (not including the three stocks in the inner gulfs of Shark Bay not addressed in this study) – Shark Bay Oceanic, West Coast and South Coast stocks (Figure 1). These stocks and their spatial boundaries are based on bioregions defined primarily from information on environmental characteristics rather than from knowledge of species-specific population structure and connectivity (Gaughan and Santoro 2021).

**FIGURE 1.**
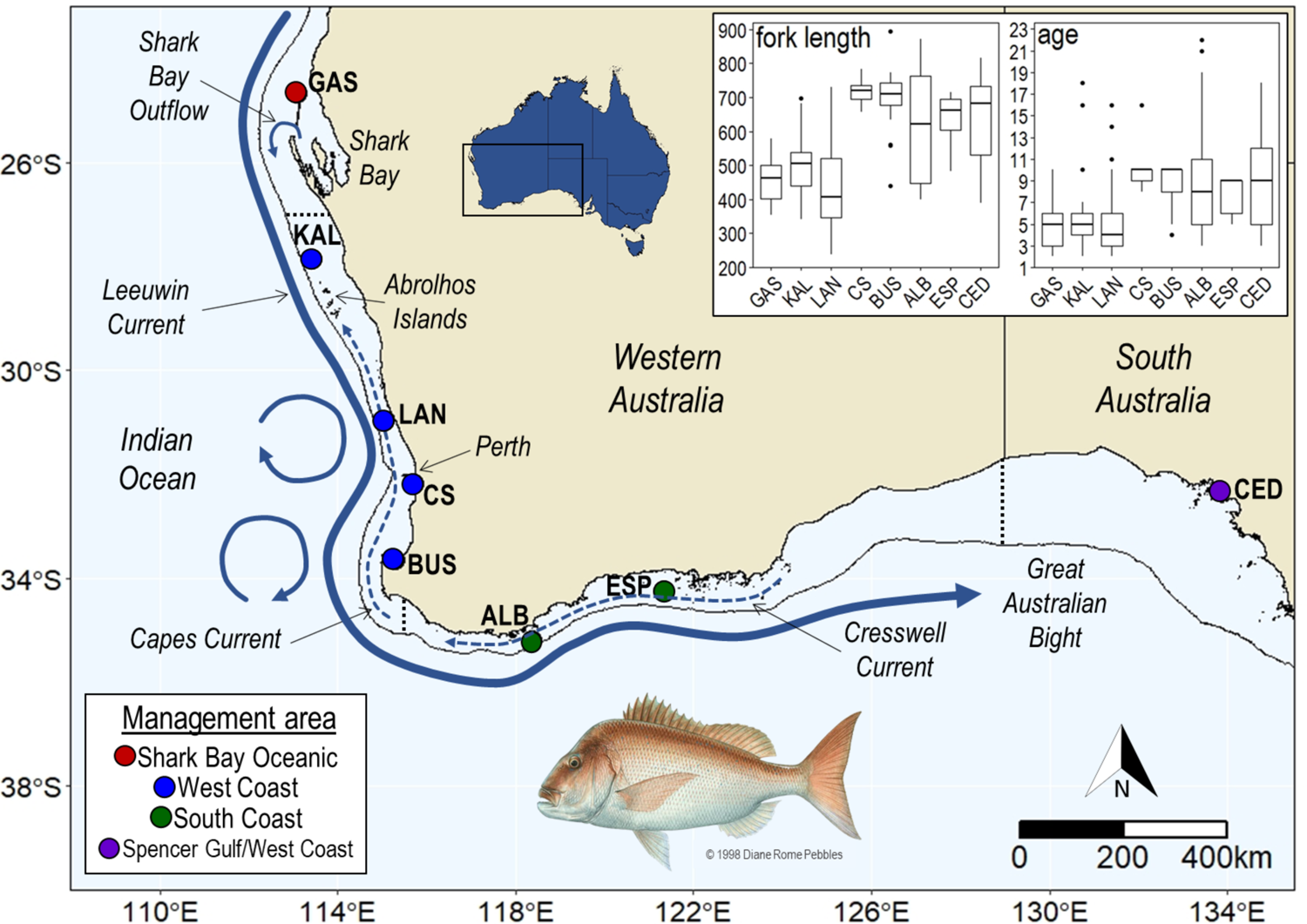
Map of the study region showing the eight sampling sites, including the comparative site in South Australia (SA), the three current management areas for snapper in Western Australia (WA): Shark Bay Oceanic (red), West Coast (blue) and South Coast (green), as well as the westernmost management area in SA: Spencer Gulf/West Coast (purple). Management area boundaries are indicated with dotted black lines. The inset plots summarise the lengths (in mm) and ages (in years) of sampled snapper (exluding the temporal samples from CS and ESP). Sampling sites in WA: GAS, Gascoyne; KAL, Kalbarri; LAN, Lancelin; CS, Cockburn Sound; BUS, Busselton; ALB, Albany; ESP, Esperance. Sampling site in SA: CED, Ceduna.

The particular reproductive characteristics and movement behaviours of snapper as well as the dominant patterns of oceanographic circulation along the WA coast are likely important in shaping stock structure and connectivity in the species. The dispersal potential of snapper is expected to be high, particularly during the pelagic and sub-adult stages, facilitating population connectivity. Snapper are multiple batch, broadcast spawners, with eggs hatching after 1-2 days and the pelagic larval stage lasting for ∼17-30 days (Francis 1994, Fowler and Jennings 2003). When eggs and larvae occur in open shelf waters they are expected to be influenced by the Leeuwin Current (LC), a boundary current that flows polewards along the west coast of WA and then eastwards along the south coast into the Great Australian Bight (Cresswell and Golding 1980; Figure 1). During the autumn and winter when snapper spawn in the north of its WA range (Shark Bay to the Albrolhos Islands; Figure 1), the LC flows most strongly and further inshore. During the late spring and early summer when snapper spawn along the lower west and south coasts of WA, eggs and larvae are expected to be influenced by a weaker LC but also by seasonal wind-driven transport systems – the northward flowing Capes Current (Pearce and Pattiaratchi 1999) and the westward flowing Cresswell Current (Cresswell and Peterson 1993, Akhir et al. 2020; Figure 1). After settlement, juvenile snapper migrate from spawning areas to sheltered inshore habitats where they reside until ∼2 years of age. During the sub-adult stage, snapper are known to migrate distances of up to over 1,000 km to offshore reefs where they become resident (Fowler et al. 2005, Hamer et al. 2011, Wakefield et al. 2011).

Other life-history and oceanographic characteristics could act to limit connectivity and dispersal in snapper in WA, leading to population structure. Although snapper spawn along the entire WA coastline, a number of embayments where large spawning aggregations have been observed are likely particularly important (Nahas et al. 2003, Wakefield 2010, Wakefield et al. 2015). These include coastal embayments of Shark Bay, Perth (Cockburn Sound, Warnbro Sound, Owen Anchorage) and Albany (King George Sound; Figure 1). Such environments are largely protected by oceanographic processes in open shelf waters and also often feature local circulation patterns that act to retain eggs and larvae (Steedman and Craig 1983, Wakefield et al. 2011). Outside of these emabyments in open shelf waters, connectivity of snapper populations could be interrupted in areas of instability of the LC where large meanders and mesoscale eddies form (Pearce and Griffiths 1991, Waite et al. 2007). With respect to older life-stages, although adult snapper are known to travel distances of hundreds of kilometres or more, most show more limited movements of spatial scales of <100 km (Moran et al. 2003, Sumpton et al. 2003, Wakefield et al. 2011, Crisafulli et al. 2019).

Current knowledge of population structure and connectivity in snapper in WA is based on tagging, otolith microchemistry and microsatellite DNA analyses. Recent tagging work on the lower west coast of WA indicated that adult movement is largely limited to spatial scales of <20 km, although this estimate is probably influenced by spatial variability in fishing effort (Crisafulli et al. 2019). Microsatellite DNA analyses have suggested that snapper in WA are characterised by an isolation by distance pattern of population structure, rather than genetically distinct sub-populations (Gardner and Chaplin 2011). Otolith microchemistry studies have reported different patterns of mixing during different life stages as well as a range of patterns of spatial differentiation using different chemical tags (Edmonds et al. 1999, Fairclough et al. 2013). Therefore, a degree of uncertainty still remains around where appropriate stock boundaries should be drawn for management and assessment purposes and whether current boundaries are appropriate. In addition, declines in snapper catches and stock depletions in parts of WA point to the need for more information about population structure and connectivity across the state (Fowler et al. 2021).

In this study, we use genome-wide markers and population genomic analyses to characterise population genetic structure and connectivity in snapper across its WA range. Our dataset also enables us to assess temporal stability in population structure as well as connectivity in snapper between WA and its nearest, eastern jurisdictional neighbor (SA). Given the species life-history traits and the oceanographic setting of WA’s coast, we predict population differentiation in snapper to be influenced by broad patterns of on-shelf oceanographic circulation. Our secondary goal is to determine whether current spatial scales of assessment and management reflect the biological units (i.e. stocks) identified with genomics, as well as their spatial boundaries. We chose a genomic approach because datasets of 1,000s of DNA markers are more suitable than previously utilised genetic datasets of 10s of markers (i.e., those used in microsatellite studies) for addressing questions about population structure and connectvity in marine populations (Grummer et al. 2019). Population genomic datasets have great power to describe these characteristics in species with very large and highly connected populations, which are features of many marine organisms (Bernatchez et al. 2017).

## 2 METHODS AND MATERIALS

### 2.1 Sampling and associated biological information

Muscle or fin-clip samples were obtained from 345 snapper landed by recreational or commercial fishermen or fisheries researchers between 2018 and 2020 at eight locations between the Gascoyne, on the central west coast of Western Australia (WA), and Ceduna, on the west coast of South Australia (SA; see Figure 1, Tables 1 and S1). These sampling locations cover the majority of fishing activity occurring in our focal jurisdiction WA, and include commercial hotspots (Kalbarri and Lancelin) and recreational hotspots near population centres (Cockburn Sound, Busselton and Albany; Gaughan and Santoro 2021).

**TABLE 1.** Levels of genome-wide variation for all snapper samples from the seven Western Australian locations (including the two temporal samples from Cockburn Sound and Esperance: CS14 and ESP10) and the comparative South Australian site (CED), based on 10,903 putatively neutral SNPs. *N*, sample size before and after (in parentheses) removing individuals with >20% missing data; FL, fork length; *H*o, observed heterozygosity; *H*e, expected heterozygosity; π, nucleotide diversity; *F*_IS_, inbreeding coefficient. No *F*_IS_ value deviated significantly from expectation (α = 0.05). Mean fork length (FL) is in mm and age in years. Following each mean FL and age is the associated standard deviation in parentheses.

| Site | *N* | Avg. FL | Avg. age | *H<sub>o</sub>* | *H<sub>e</sub>* | $\pi$ | *F<sub>IS</sub>* |
| --- | --- | --- | --- | --- | --- | --- | --- |
| Gascoyne (GAS) | 40 (39) | 457.4 (61.5) | 5.1 (2.0) | 0.181 | 0.185 | 0.188 | 0.029 |
| Kalbarri (KAL) | 40 (40) | 497.3 (84.8) | 5.7 (3.5) | 0.182 | 0.186 | 0.188 | 0.025 |
| Lancelin (LAN) | 37 (37) | 441.6 (130.8) | 5.1 (3.3) | 0.184 | 0.186 | 0.189 | 0.020 |
| Cockburn Sound (CS) | 40 (39) | 713.5 (30.0) | 9.7 (0.6) | 0.186 | 0.187 | 0.189 | 0.012 |
| Cockburn Sound 2014 (CS14) | 30 (29) | 794.9 (42.7) | N/A | 0.180 | 0.184 | 0.187 | 0.016 |
| Busselton (BUS) | 40 (40) | 702.8 (72.1) | 9.0 (1.5) | 0.180 | 0.185 | 0.187 | 0.030 |
| Albany (ALB) | 40 (39) | 611.1 (163.0) | 8.9 (5.5) | 0.189 | 0.188 | 0.191 | 0.008 |
| Esperance (ESP) | 13 (13) | 631.8 (92.9) | 7.6 (1.9) | 0.187 | 0.182 | 0.189 | 0.007 |
| Esperance 2010 (ESP10) | 28 (28) | N/A | N/A | 0.184 | 0.186 | 0.189 | 0.015 |
| Ceduna (CED) | 37 (37) | 634.2 (127.2) | 8.7 (4.1) | 0.188 | 0.187 | 0.190 | 0.005 |

Tissues from ∼40 individuals per location were preserved in 100% ethanol and stored at −20°C until DNA extraction. Where possible, sagittal otoliths and biological data including total and fork length, sex and reproductive stage were obtained for each sampled individual (see Tables 1 and S1). Additional tissue samples were obtained from snapper landed off Esperance and in Cockburn Sound in 2010 and 2014, respectively, to investigate the temporal stability of trends in population structure. The Cockburn Sound samples were collected by fisheries researchers as part of a fisheries independent survey, while the Esperance samples were collected for a former population genetics project (Gardner and Chaplin 2011, Gardner et al. 2014, Gardner et al. 2017).

### 2.2 DNA extraction, genomic library preparation and sequencing

Genomic DNA was isolated from each sample using a modified salting-out protocol (Sunnucks and Hales 1996). Quality control of DNA extracts was carried out with NanoDrop and gel electrophoresis. Extracts were quantified with Qubit and diluted to ∼10-15 ng/μL. Double-digest restriction site-associated DNA (ddRAD) libraries were constructed following a protocol modified from Peterson et al. (2012), as detailed in Brauer et al. (2016). For each sample, 200 ng of genomic DNA was digested using the restriction enzymes Sbfl-HF and Msel (New England Biolabs). One of 96 unique 6 bp barcodes was ligated to each sample before pooling libraries into groups of 12 samples. DNA fragments between 300 and 800 bp were selected from each pool using a Pippin Prep (Sage Science). Each pool was then amplified in three 25 μL reactions to reduce PCR artefact bias. Following PCR, the three reactions were combined, and the size distribution of the products examined using a 2,100 Bioanalyser (Agilent Technologies) and quantified using Qubit. Aliquots of equal concentrations were then taken from each pool and combined to form one pool of 96 samples. Pools were sequenced on an Illumina HiSeq 4,000 (150 bp paired end) at Novogene (Hong Kong). Six replicates were included in each pool of 96 samples so that sequencing and genotyping errors could be quantified.

### 2.3 Bioinformatics

Raw sequence reads were processed to generate a high-quality SNP dataset using similar bioinformatic procedures as detailed elsewhere (e.g. Sandoval-Castillo et al. 2018), but with the assistance of a reference genome for snapper. Specifically, the quality of raw sequence data was checked using FastQC before being demultiplexed with the process_radtags module from STACKS 2.0 (Catchen et al. 2013). Barcodes, restriction sites and RAD tags were then trimmed from sequence reads using TRIMMOMATIC (Bolger et al. 2014). Trimmed sequence reads were then aligned to a high-quality snapper reference genome (Catanach et al. 2019) using BOWTIE 2 (Langmead and Salzberg 2012). The SNPs were subsequently called using BCFTOOLS (Narasimhan et al. 2016). The resulting dataset was initially filtered using VCFTOOLS (Danecek et al. 2011) to retain only bi-allelic SNPs present in at least 80% of individuals in all populations with a minimum minor allele frequency of 0.03. Also using VCFTOOLS, further filtering was carried out to remove indels, individuals with more than 20% missing data, SNPs with low and extremely high coverage, SNPs with low mapping quality, SNPs not in Hardy-Weinberg Equilibrium (HWE) and physically linked SNPs.

### 2.4 Categorising putatively neutral SNPs

The Bayesian method in BAYESCAN 3.0 (Foll and Gaggiotti 2008) was used to identify candidate SNPs putatively under selection. The software was run with 20 pilot runs, each with 5,000 iterations, followed by 100,000 iterations with a burn-in length of 50,000 iterations. The outlier SNPs were identified using a 5% false discovery rate with a prior odd of 10 and were subsequently removed from the dataset to produce a putatively neutral one. The study of the role of natural selection on snapper populations using outlier SNPs and genomic regions associated with environmental variation is the topic of a separate and ongoing investigation (Brauer et al. unpublished).

### 2.5 Genetic diversity, population differentiation and clustering analyses

The genetic diversity statistics observed heterozygosity (*H*_O_), expected heterozygosity (*H*_E_) and nucleotide diversity (π), were calculated in STACKS 2 (Rochette et al. 2019). Pairwise *F*_ST_ and population specific *F*_IS_ values were calculated in ARLEQUIN 3.5 (Excoffier and Lischer 2010), with significance assessed with 1,000 permutations. *F*_ST_ associated *p*-values were subsequently corrected for multiple comparisons through the false discovery rate (FDR) method using the *p.adjust* function in the R package BASE 4.0.3 (Team 2021). Global *F*_ST_ was determined using the *basic.stats* function in the R package HIERFSTAT 0.5-10 (Goudet et al. 2015). The model-free approach, Principal Components Analysis (PCA), was then carried out using VEGAN 2.5-6 (Oksanen et al. 2018) in R. Missing genotypes (∼0.6% of data matrix) were assigned with the most common genotype at that locus. Population structure was further assessed with the maximum likelihood approach implemented in ADMIXTURE 1.3 (Alexander et al. 2009, Alexander and Lange 2011). The cross-validation procedure in ADMIXTURE was employed to determine the most likely K value. To do this, a 5-fold cross-validation was performed for K values 1-8. Graphical representation of population assignments was performed with GGPLOT2 3.3.3 (Wickham 2016) in R. To assess temporal stability in patterns of population structure, the ADMIXTURE analysis was carried out again including the two temporal samples and membership to any identified clusters was compared between sampling periods.

### 2.6 Isolation by coastal distance

We tested for a signal of isolation by distance (IBD) across the sampling range by assessing the relationship between coastal distance and linearized *F*_ST_ (*F*_ST_/(1-*F*_ST_)) with a Mantel test (9,999 permutations) in GENALEX 6.5 (Peakall and Smouse 2012). Distances between sampling locations were estimated as the shortest distance between sites following the coastline using the *viamaris* function in MELFUR 0.9 (https://github.com/pygmyperch/melfuR). Mean effective dispersal distance was then estimated from the slope of the IBD relationship using the theoretical model of Kinlan and Gaines (2003), an extension of Palumbi (2003): mean dispersal distance = 0.0016(IBD slope)^-1.0001^. The resulting estimate equates to the mean dispersal distance required to produce the obersved IBD slope under model assumptions about parameters such as effective population size and population density. This dispersal model allows for inferences about the spatial scales over which individuals successfully disperse and establish on a per generational basis and therefore provides information on demographic connectivity, a concept important in fisheries management (Palumbi 2003). Of further relevance to fisheries management is that IBD slopes, unlike *F*_ST_, are not affected by rare dispersal events but strongly reflect dispersal over the most proximate generations and therefore over ecologically-significant time scales (Rousset 1997).

### 2.7 Connectivity barriers

We tested for putative barriers to dispersal using the piecewise regression approach implemented in Robinet et al. (2020). Briefly, the individual ancestry proportions generated by ADMIXTURE for a K of 2 (presented in Figure 2c) were used to explore the presence of breaks in the admixture gradient across samples with significant WA ancestry (i.e., between the Gascoyne and Esperance). A piecewise regression of mean SA ancestry (i.e., ancestry characteristic of the SA site Ceduna) as a function of distance from the last sample with individuals exhibiting significant WA ancestry (i.e. Esperance) was performed using the R script Introgression_breaks.R (https://github.com/tonyrobinet/introgression). A barrier to dispersal was considered present when the piecewise model provided a significant reduction in residual sum of squares relative to the simple regression model.

**FIGURE 2.**
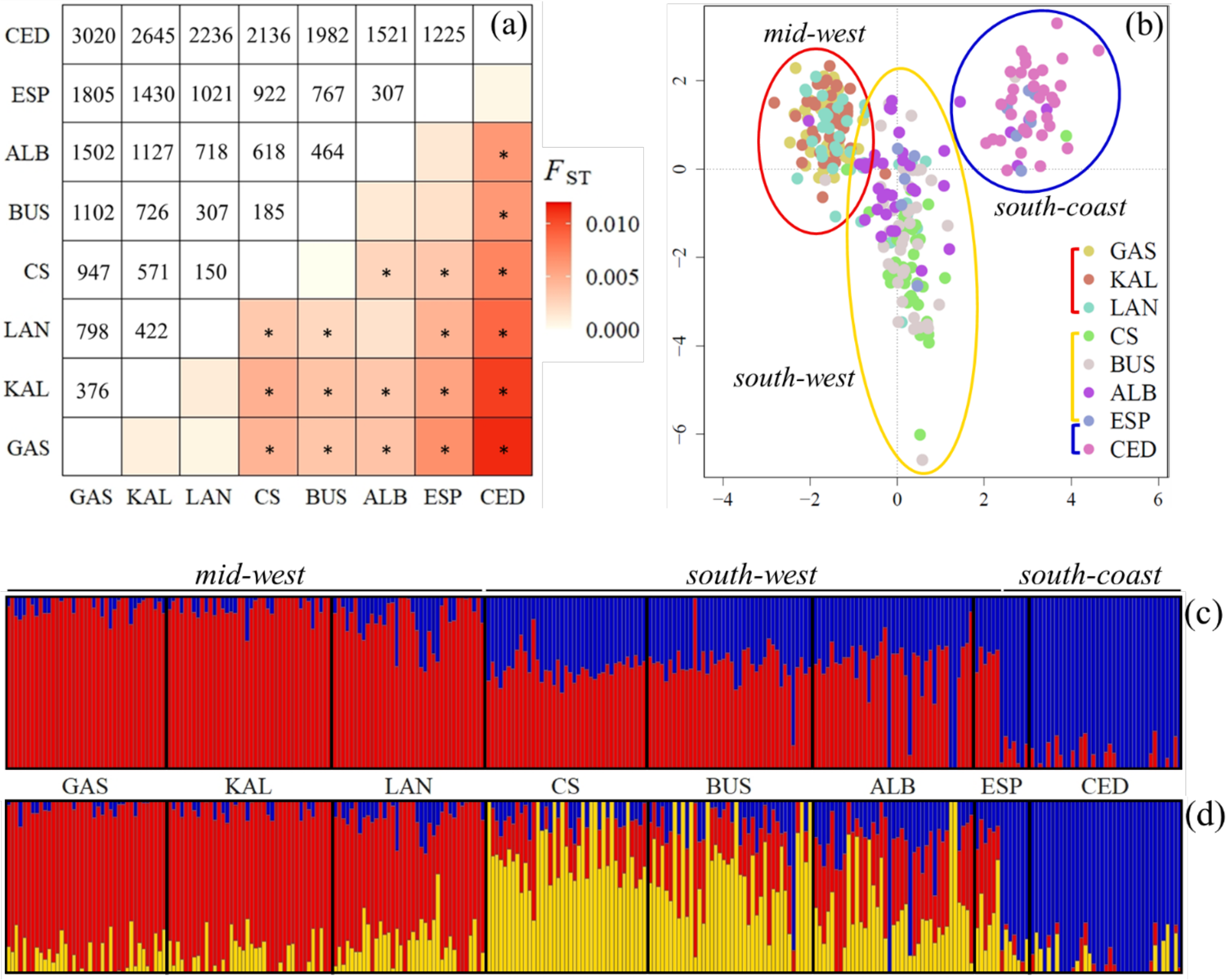
Pairwise estimates of genetic differentiation (a) and broad-scale genetic clusters (b-d) for snapper between the central west coast of Western Australia (GAS) and the west coast of South Australia (CED). These results are based on the 10,903 neutral SNPs and exclude the two temporal samples. More specifically: (a) a heatmap of pairwise *F*_ST_ values (below diagonal) and the coastal distances between sampled sites (km; above diagonal), with the asterisks indicating *F*_ST_ values that remained significant (α = 0.05) after correction for multiple comparisons via the FDR estimation. (b) Principal component analysis (PCA) of genetic differentiation, with principal component 1 (PC1: 0.87% of variance) against principal component 2 (PC2: 0.59% of variance) and each point representing an individual, colour coded by sampling location, and the circles indicating the three identified genetic clusters – the mid-west (GAS, KAL, LAN), the south-west (CS, BUS, ALB) and the south-coast (CED). Note that the ESP sample spans the south-west and south-coast groups. Bar plots of the ADMIXTURE clustering analysis results for (c) K = 2 and (d) K = 3. Labels above the plots represent the three groups identified with the PCA. Individuals are represented by the vertical bars and each individual is coloured according to its probability of membership to each of the three clusters, which are represented by red, yellow and blue.

### 2.8 Local-scale structure

To investigate local-scale patterns of gene flow and further explore the impact of coastal distance on genetic structure, we assessed the genetic similarity between individuals at increasing geographic distances with spatial autocorrelation analyses in GENALEX 6.5 (Smouse and Peakall 1999, Peakall and Smouse 2012). Spatial autocorrelation analysis can provide higher resolution information on current patterns of gene flow than evolutionary estimators like *F*_ST_ (Peakall et al. 2003). Specifically, the analysis can uncover IBD signals over smaller scales and demarcate ecologically important genetic patches. The extent of nonrandom and positive spatial autocorrelation can be inferred from the first x-intercept in the correlogram (also referred to as genetic patch size) if a significant correlation coefficient (*r*) occurs in at least one distance class (Sokal and Wartenberg 1983, Smouse and Peakall 1999). Analyses were done separately for the two main WA groups identified with the clustering and piecewise regression analyses (i.e., the mid-west and south-west groups; see Results). Distance classes were chosen such that sample sizes per class were adequate and approximately equal. *R* values were also calculated for all eight samples separately to assess within-location autocorrelation. The significance of each *r* value was determined with 1,000 bootstraps, while 95% CIs around the null hypothesis of randomly distributed genotypes were determined with 1,000 permutations. To reduce the possibility of overinterpreting the correlograms, we only considered a value of *r* to be significant if it fell outside of the CIs around the null hypothesis of zero correlation and if its error bars did not cross the x-axis.

## 3 RESULTS

### 3.1 SNP genotyping

A total of 7,342,804 raw SNPs were characterised. After completing all filtering steps and removing candidate adaptive SNPs, our final ddRADseq dataset comprised 10,903 putatively neutral SNPs (details in Table S2). Four of the 345 sequenced samples were removed from the dataset due to having >20% missing data, leaving 341 snapper for subsequent analyses (Table 1). The 341 samples had an average of 1.1% missing data (range: 0.009-14.1%).

### 3.2 Genetic diversity, population differentiation and clustering analyses

Levels of genetic diversity were very similar across sites (Table 1). Expected heterozygosity (*H*_E_) ranged from 0.182 to 0.188 and observed heterozygosity (*H*_O_) ranged from 0.180 to 0.189. Values of the population specific inbreeding coefficient (*F*_IS_) were close to zero for all sites (range: 0.005-0.03) and none deviated significantly from expectation. Genetic differentiation (*F*_ST_) between pairs of sampled sites was nil to low and ranged from 0 to 0.011, with CED being the most differentiated sample (Figure 2a). Despite the low differentiation, 18 of the 28 pairwise site comparisons remained significant after correction for multiple comparisons via FDR estimation. Global *F*_ST_ for the species between the central west-coast of WA (GAS) and the west-coast of SA (CED) was 0.003.

The PCA indicated the presence of three geographically distinct groups across the study region, referred to herein as the mid-west (GAS, KAL, LAN), south-west (CS, BUS, ALB) and south-coast (CED; Figure 2b). The ESP sample did not cluster predominately with any one group but spanned across the south-west and south-coast. Differentiation between the groups varied, with greater distinction observed between the south-coast and the south-west and mid-west groups than between the mid-west and south-west groups. Of the two WA groups, the mid-west was more tightly clustered than the south-west, suggesting greater homogeneity in the former.

The ADMIXTURE analysis suggested that the most probable number of genetic clusters in the dataset was two (i.e., K = 2). These clusters loosely corresponded to the samples between GAS and ALB (plus ∼ half of the ESP sample) and the South Australia (SA) sample (CED; plus the other half of the ESP sample; Figure 2c). However, inspection of the *q* values for K = 2 indicated the presence of two weakly differentiated groups within the larger cluster with clearly distinct ancestry proportions. These groups corresponded with the mid-west and south-west clusters identified with the PCA. The ancestry proportions for K = 3 distinguished these groups and provided further information on population structure across the study region (Figure 2d). For example, they indicated greater homogeneity in ancestry proportions within the mid-west than the south-west group, suggesting that biologically relevant fine-scale structure may occur within the latter.

Comparing the two temporal CS and ESP samples, membership to the three groups identified with ADMIXTURE indicated temporal stability in genetic structure at the former location only (Figure 3). Compared with the ESP sample, ESP10 had higher membership to the south-coast group (Wilcoxon test: *p*-value < 0.0005) and lower membership to the mid-west and south-west groups (Wilcoxon test: *p*-value = 0.001 and 0.02, respectively).

**FIGURE 3.**
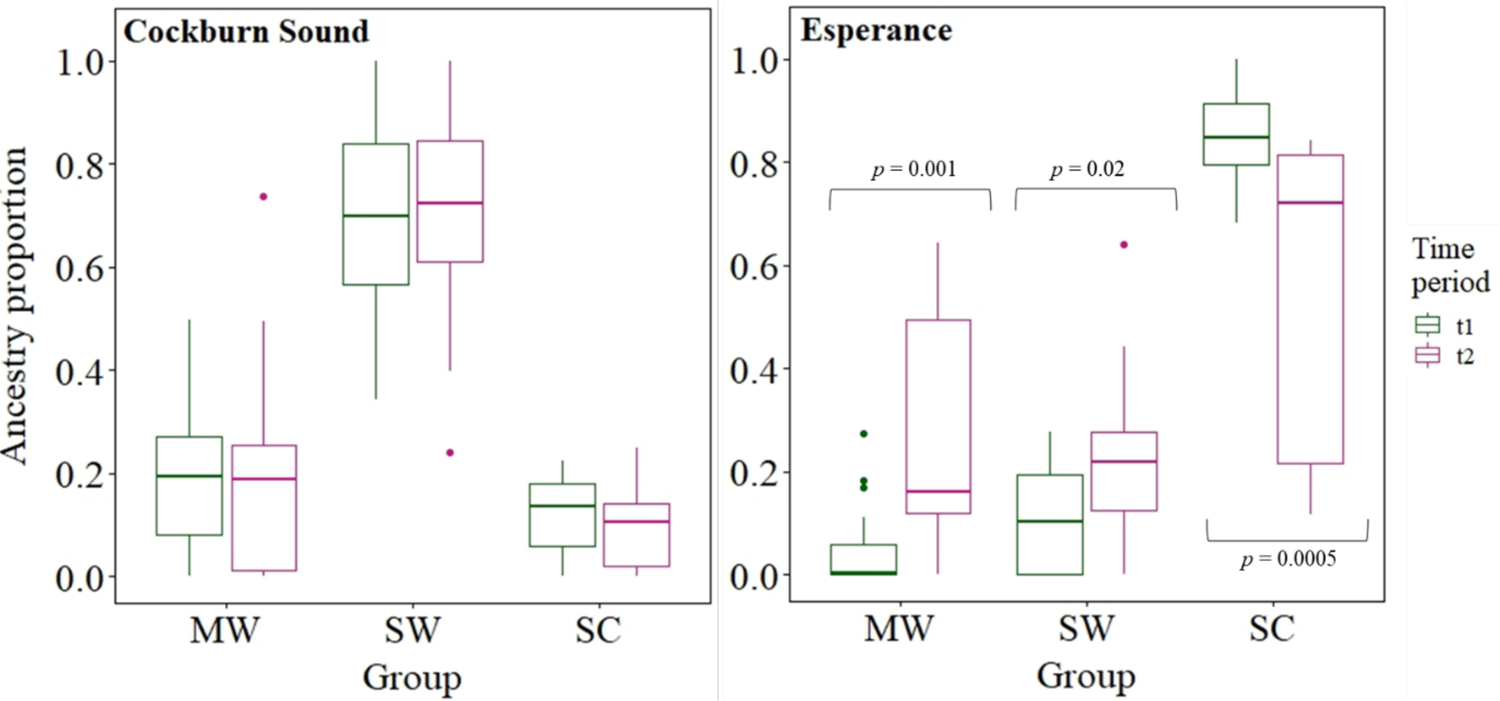
Summary of membership to the three identified groups (i.e., MW, mid-west; SW, south-west; SC, south-coast) for the temporally separated snapper samples from CS and ESP. Ancestry membership proportions did not significantly differ between the two CS samples, but differed for all three comparisons for the ESP samples. Ancestry proportions are from the ADMIXTURE results for K = 3. For CS, t1 = 2014 and t2 = 2018, while for ESP, t1 = 2010 and t2 = 2019/2020.

### 3.3 Isolation by coastal distance

We detected highly significant IBD across the sampling range (r = 0.86, *p*-value < 0.001; Figure 4). This analysis indicated that spatial distance explains 74% of the variation in linearized *F*_ST,_ accounting for a substantial amount of the population genetic differentiation inferred across the sampling range. Mean effective dispersal distance per generation was estimated as 400 km from an IBD slope of 4E-06 (i.e., *F*_ST_ = 0.004 per 1,000 km). This suggests that demographic connectivity between locations separated by distances greater than 400 km may be limited. Our estimate of 400 km is within the range of those made for other marine fish (Kinlan and Gaines 2003).

**FIGURE 4.**
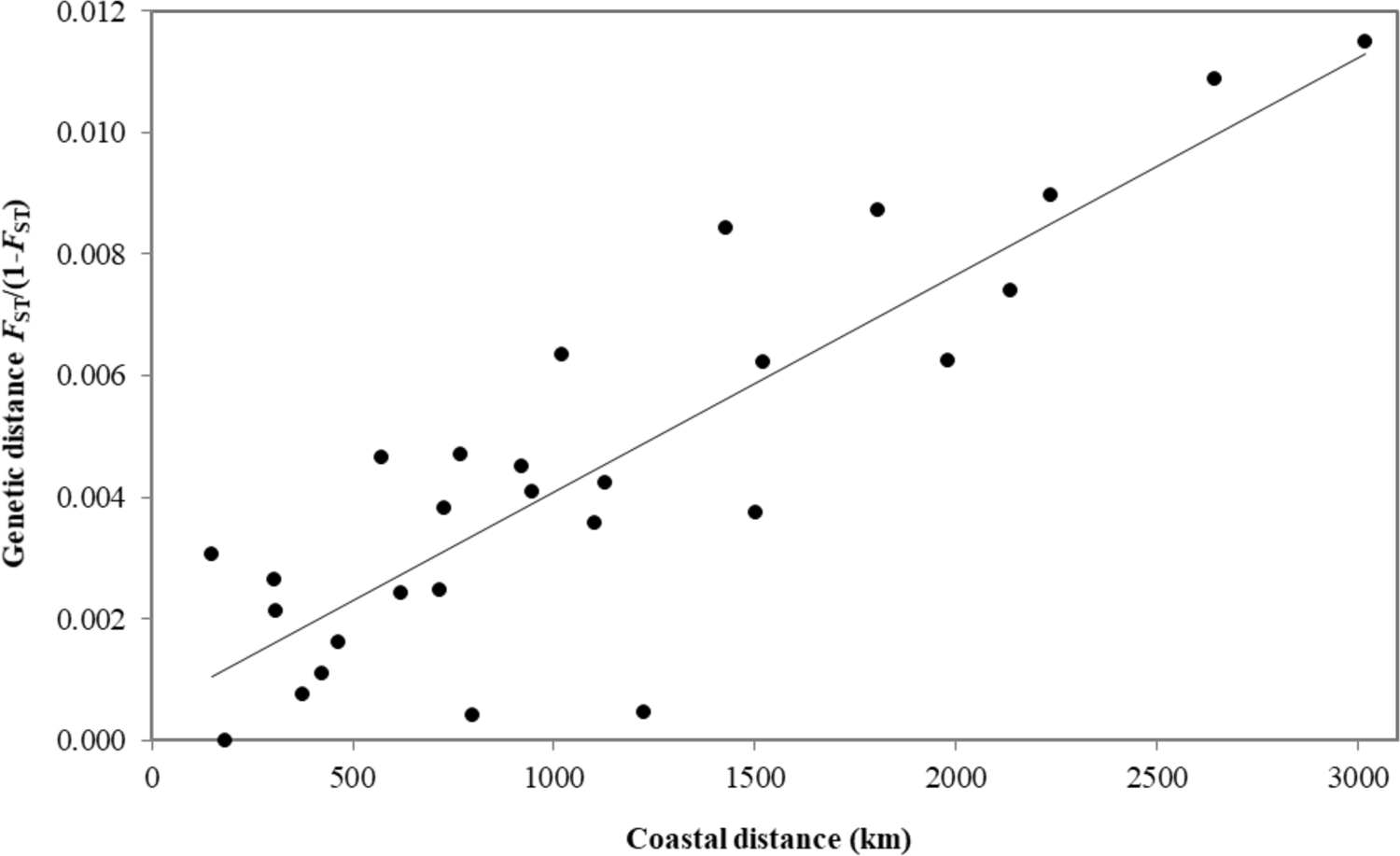
Relationship between coastal distance and genetic distance (linearized *F*_ST_) for snapper between the central west coast of Western Australia (GAS) and the west coast of South Australia (CED; Mantel test: r = 0.86, *p*-value < 0.001).

### 3.4 Barriers to connectivity

The piecewise regression analysis identified a significant break point in the ancestry gradient between CS and LAN, despite these sites being separated by only ∼150 km (Figure 5). Although the simple linear regression model revealed a strong negative relationship between distance and mean SA ancestry (adj. r^2^ = 0.67, *p*-value = 0.03), the piecewise regression model was a significantly better fit to the data (piecewise model: adj. r^2^ = 0.98, *p*-value = 0.01; ANOVA for model comparison: *p*-value < 0.001). These results suggest that the genetic structure observed along the WA coast is not merely due to IBD but is also a reflection of a connectivity barrier, an interpretation consistent with results of the clustering analyses.

**FIGURE 5.**
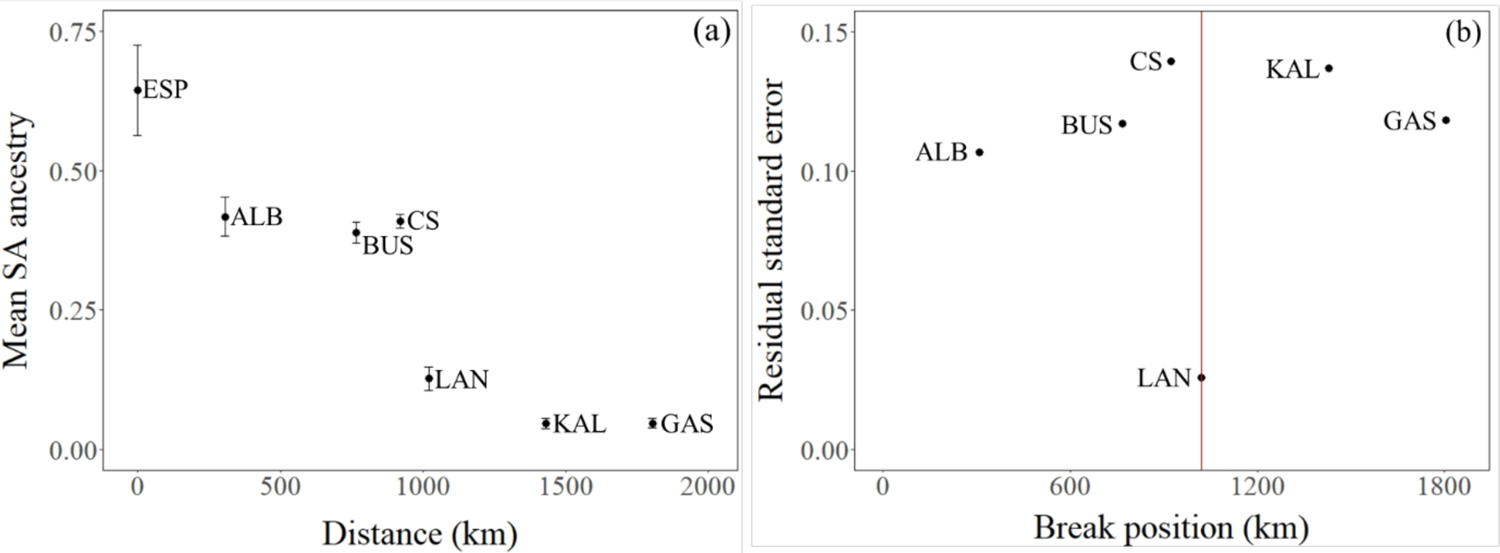
Analysis of barriers to connectivity in snapper along the Western Australia (WA) coast. (a) Mean proportion of South Australian (SA) ancestry (i.e., ancestry characteristic of the blue group CED in Figure 2c) across the WA samples (± standard error), determined from the ADMIXTURE results for K = 2, as a function of distance from ESP, and (b) results of the piecewise regression analysis showing residual standard error as a function of distance from ESP.

### 3.5 Local-scale structure

The spatial autocorrelation analyses indicated that across the sampling range, genetic autocorrelation occurs between individuals sampled at the same site (Figure 6c), a result suggestive of recruitment to the local subpopulation. Within site spatial autocorrelation was highest at GAS and KAL in the mid-west and at ALB in the south-west (Figure 6c). Variation around sample-specific *r* values was highest for KAL, ALB and ESP. These samples were the result of the greatest number of fishing days that were separated by the greatest stretches of time (Table S1). This could perhaps indicate that cohesion among different groups of genetically alike individuals may occur in snapper across this region, but such a possibility would need to be verified with larger samples than those available here.

**FIGURE 6.**
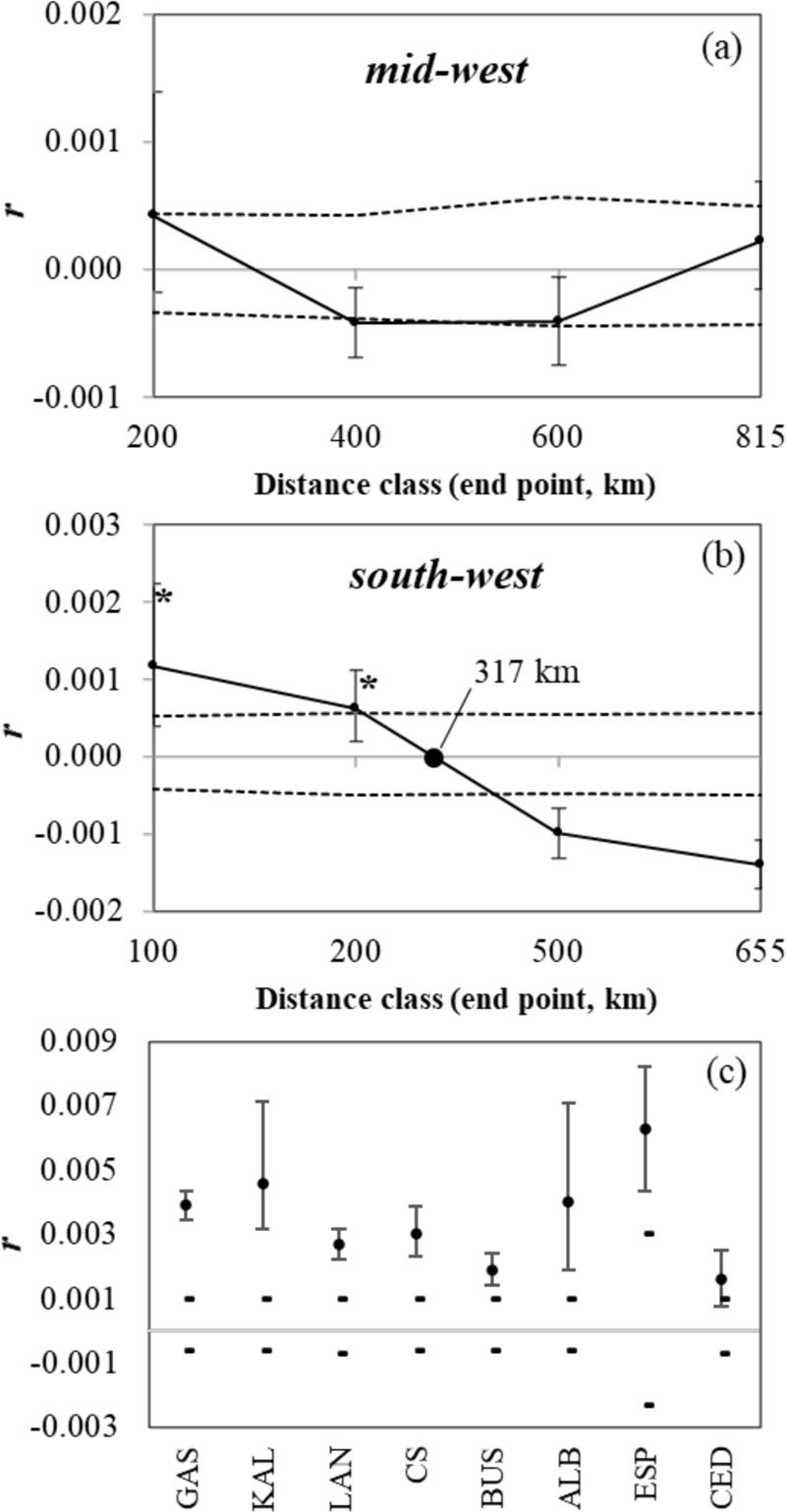
Spatial autocorrelation analyses of snapper samples from the (a) mid-west, (b) south-west and (c) each of the eight samples separately. These analyses exclude the two temporal samples and are based on 10,903 neutral SNPs. Values of *r* represent genetic autocorrelation coefficients, the dashed lines represent 95% CIs around the null hypothesis of randomly distributed genotypes, determined with 1,000 permutations, and the error bars represent 95% CIs around *r* values, as determined by 1,000 bootstraps. In the correlograms, the first x-intercept following significantly positive values of *r* (denoted by asterisks) indicates the extent of positive spatial autocorrelation or the genetic patch size.

Our spatial autocorrelation correlograms indicated that local-scale genetic structure occurs within the south-west but not the mid-west (Figure 6a-b), a result concordant with observations made from the PCA and ADMIXTURE plots. In the mid-west, genetic autocorrelation occurred only among individuals landed at the same site, suggesting that it is largely a homogenous group (Figure 6a, c). In contrast, in the south-west, significant positive spatial autocorrelation also occurred in the 100-200 km distance class (Figure 6b). The shape of the correlogram was of the ‘long-distance cline’ type (Diniz-Filho and De Campos Telles 2002, Östman et al. 2017), suggesting that IBD occurs across the south-west. The south-west had an x-intercept and therefore a genetic patch size of ∼300 km, indicating the scale over which high levels of gene flow occur. This estimate also suggests that in the south-west, individuals separated by >300 km are less genetically similar than expected at random.

## 4 DISCUSSION

This is the first study to investigate population structure and connectivity in the fishery important snapper in Australia using a population genomics approach. We detected spatially variable population genetic structure and levels of connectivity in snapper across ∼3,000 km of coastline. Although our results indicated that snapper across Western Australia (WA) are characterised by weak genetic structure, we found evidence for stock discontinuities, dispersal limits and local-scale structure. Our results improve upon prior understanding of population structure and connectivity in snapper across its WA range based on tagging (Wakefield et al. 2011, Crisafulli et al. 2019), otolith microchemistry (Edmonds et al. 1999, Fairclough et al. 2013) and microsatellite analyses (Gardner and Chaplin 2011). The variable patterns of population dynamics inferred along the WA coast appear influenced by a combination of species life-history, spawning behaviour and on-shelf oceanographic processes. Our results partly contrast with the spatial scales of assessment and management currently used for snapper in WA, highlighting the need for a review. Our study supports the value of population genomic surveys to inform the management of weakly-structured and wide-ranging marine fishery resources.

### 4.1 Broad-scale structure

We uncovered two broad-scale patterns of population genetic structure in snapper across WA with our neutral SNP dataset. Firstly, strong isolation by distance (IBD) was detected between the central west-coast of WA and the west-coast of South Australia (SA), indicating that dispersal is limited by coastal distance and probably primarily occurs in a stepping-stone fashion. This suggests that the extent of effective dispersal per generation across all life-stages is limited by distance. The IBD relationship indicated that this distance is likely ∼400 km on average. The WA data on adult movement suggests that most remain within spatial scales of ∼100 km (Moran et al. 2003, Wakefield et al. 2011, Crisafulli et al. 2019).

Hypothetically, the longer-range dispersal required to produce the observed IBD relationship along the WA coast may occur at the larval and/or sub-adult stage. Indeed, otolith microchemistry work on snapper has suggested that movement of individuals post recruitment may be greatest at the sub-adult stage (Fowler et al. 2005, Hamer et al. 2011, Fairclough et al. 2013, Fowler et al. 2017). Previous work with microsatellite markers and otolith microchemistry has also suggested that IBD occurs in snapper across WA (Edmonds et al. 1999, Gardner and Chaplin 2011, Fairclough et al. 2013). IBD has also been detected in a number of other broadcast spawning marine species within our study range, including Roe’s abalone (Hancock 2000), WA dhufish (Berry et al. 2012) and saucer scallop (Kangas et al. 2019).

Broad-scale IBD signals do not convey information on whether genetic discontinuities exist or where they are located. The clustering and connectivity barriers analyses allowed us to detect two genetic discontinuities, which lie on the lower west coast and in the southeast of the study range. Although Gardner and Chaplin (2011) also detected the discontinuity in the southeast using microsatellite markers, our study is the first to report a discontinuity for snapper on the lower west coast. Otolith microchemistry work with snapper along the west coast of WA indicated differentiation between snapper north and south of the Abrolhos Islands, located on the mid-west coast (Figure 1; Edmonds et al. 1999). However, that study lacked samples from locations between the Abrolhos Islands and Cockburn Sound so could not have detected a discontinuity just south of Lancelin. Additionally, although another otolith microchemistry study detected differentiation in snapper along the west coast of WA, a clear stock discontinuity was not detected between Cockburn Sound and Lancelin (Fairclough et al. 2013). The discontinuities uncovered here allowed us to delineate three broad-scale snapper stocks between the Shark Bay region and the west coast of SA (i.e., the mid-west, south-west and south-coast stocks; see Figure 2).

Generally, the stock discontinuities reported in this study lie between important spawning sites across WA and SA where the largest aggregations of spawning snapper have been observed – the Shark Bay region, Cockburn Sound (including the adjacent embayments Owen Anchorage and Warnbro Sound) and the SA gulfs. Therefore, these discontinuities may indicate the importance of these spawning areas in maintaining the three stocks. The locations of the genetic discontinuities may reflect the spatial extent of both recruitment from spawning events at these breeding sites as well as effective dispersal of individuals post-recruitment. These processes are probably largely influenced by physical factors like oceanography and coastal topography as well as biological factors like population density and resource availability.

In the Shark Bay region, snapper spawn between May and September (Wakefield et al. 2015) when the southward flowing Leeuwin Current (LC) is strongest (Cresswell and Golding 1980, Godfrey and Ridgway 1985). The southwards dispersal of pelagic life stages (PLS) produced around Shark Bay therefore has the potential to be extensive, especially considering the pelagic larval duration of snapper (∼18-30 days). The transport of PLS from oceanic areas of Shark Bay is likely further facilitated by the Shark Bay Outflow, which exits the embayment to the north and west before travelling southwards (Figure 1; Hetzel et al. 2018). Particle tracking modelling assessing dispersal of WA shelf waters suggests that PLS produced around Shark Bay could be transported across ∼5° of latitude (Feng et al. 2010). In contrast to Shark Bay, snapper spawn in Cockburn Sound between October and December when the LC is weakest and the northward flowing Capes Current (CC) is present (Figure 1; Pearce and Pattiaratchi 1999). Particle tracking modelling suggests that PLS generated at the latitude of Cockburn Sound would mostly be retained locally, with some small scale northward and southward transport also occurring (Feng et al. 2010). However, the magnitude of dispersal of PLS of snapper from Cockburn Sound is probably limited because of wind-driven gyres within the embayment (Steedman and Craig 1983), which have been shown to facilitate the retention of eggs and larvae (Wakefield 2010).

The discontinuity between the mid-west and south-west stocks is remarkable considering that it lies between our two most proximate sampling locations (Lancelin and Cockburn Sound), separated by only 150 km. The discontinuity lies between both inshore and offshore ecological bioregions (inshore: Southwest Shelf Transition and Southwest Shelf Province, offshore: Central Western Transition and Southwest Transition) delineated from patterns of species distributions and environmental features (Commonwealth of Australia 2006). The boundary between these bioregions coincides with patterns in the distribution of invertebrates (Kott 1952, O’Hara and Poore 2000), fishes (Hutchins 1994, Ayvazian and Hyndes 1995, Williams et al. 2001, Last et al. 2011) and seaweeds (Wernberg et al. 2013). These studies, along with ours, indicate that there may be oceanographic features in the region capable of disrupting alongshore transport of PLS. Indeed, the marine environment around Lancelin is dynamic and comprises features that could achieve this. South of the Abrolhos Islands (Figure 1) the LC strengthens, leading to its instability and the production of mesoscale features like undulations, offshoots, meanders and eddies (Pearce and Griffiths 1991). Eddies generated north of Lancelin interact with the coastline at Jurien Bay (∼30°S) where the shelf narrows, causing the offshore movement of water (Pearce and Griffiths 1991, Pattiaratchi 2006, Feng et al. 2007). Circulation patterns off Lancelin, along with the retention capability of the gyres in Cockburn Sound, may contribute to the genetic discontinuity observed for snapper between the mid-west and south-west regions.

Our results also suggest that movement of sub-adult and adult snapper between the mid-west and south-west is not substantial enough to prevent the formation of a genetic discontinuity. Although the majority of recaptures of snapper tagged in Cockburn Sound have occurred within ∼20 km, individuals have been recaptured at up to 700 km to the north and 250 km to the south (Crisafulli et al. 2019). Our results suggest that such long distance northward movements may be rare, as well as any southward movements from the mid-west. Differences in biological characteristics have been reported in snapper between the mid-west and south-west (Wakefield et al. 2015), also seen in the length and age data of our samples (see Figure 1), supporting the idea of limited movement between the two regions. These results indicate that there may be differences in resource availability and/or population density between the two areas, as well as variation in preferences of environmental conditions. Indeed, differences in environmental features exist between the ecological bioregions around the snapper stock discontinuity, including sediment type, geology, temperature, waves and dissolved oxygen (Thackway 1998, Heap et al. 2005, Lyne et al. 2005). Identification and analysis of genomic regions associated with environmental variation (e.g. Sandoval-Castillo et al. 2018, Grummer et al. 2019), may shed light on divergent ecological adaptations of snapper in each bioregion.

In SA, snapper spawn between November and February. Transport of PLS from the two SA gulfs is likely negligible because the thermal fronts present at their entrances during the summer largely prevent gulf-shelf exchanges (Bruce and Short 1990, Vaz et al. 1990, Petrusevics 1993). For example, in *Nerita atramentosa* (an intertidal snail with a pelagic larval duration of around four months) the percentage of larvae released during summer in the two SA gulfs that did not reach the boundary current ranged from 70 to 100% (Teske et al. 2015). Hypothetically, long-distance dispersal may occur during post-recruitment stages in this region. Our results suggest that the spatial scales of dispersal from SA occurring in a westward direction are remarkably large. Individuals with ancestry characteristic of our SA sample were detected as far west as Busselton, located on the lower west coast of WA, almost 2,000 km from our SA site. This finding agrees with the work of Fowler et al. (2005) showing that very long distance dispersal occurs in snapper in SA, which they suggested takes place during the sub-adult stage. The position of the genetic discontinuity in the south-east of our study region may in part reflect the spatial extent of density-dependent dispersal from SA (relative to dispersal from areas to its west), which in turn likely depends on year-to-year recruitment strength and local population densities. Indeed, interannual recruitment variability can be substantial in snapper and is considered to majorly influence population dynamics (Francis 1993, Hamer and Jenkins 2004, Fowler and McGlennon 2011, Wakefield et al. 2015).

### 4.2 Local-scale structure

The strength of an IBD relationship may be greater in some areas and weaker in others. The spatial autocorrelation analyses allowed us to investigate whether local-scale patterns of genetic structure in WA mirror the broad-scale signal of IBD. Our results indicated differences in spatial genetic structure between the two WA stocks, an outcome consistent with observations made from the clustering analyses. In the mid-west, we only detected significant genetic autocorrelation among individuals from the same location. This suggests that, although a degree of cohesion between genetically similar individuals occurs within this region, connectivity and genetic homogeneity occur over large spatial scales, likely facilitated by the LC. The mid-west therefore deviates from the broad-scale IBD pattern and constitutes a well-mixed stock of snapper.

In the south-west, we detected a pattern of spatial autocorrelation consistent with IBD. Positive spatial autocorrelation occurred at distances up to ∼300 km, suggesting that demographic connectivity between our sites on the western and southern coastlines of the region is probably limited and may occur in a stepping-stone fashion and therefore indirectly over generations. King George Sound in Albany is a known important spawning area for snapper along the south coast of WA (Wakefield et al. 2015). It is possible that this embayment, along with a number of smaller adjacent embayments, are most important for supplying parts of the southern coastline of the south-west (Potter et al. 1990, Neira and Potter 1992). Supporting this idea, otolith microchemistry work with snapper suggested limited connectivity across the south-west corner of WA (Fairclough et al. 2013). The IBD pattern in the south-west may result from a weak LC during the spawning period of snapper in the region and the location of important spawning sites (i.e., protected embayments). Overall, our spatial autocorrelation analyses suggest that the influence of oceanographic processes on the transport of PLS may be particularly important in shaping patterns of genetic structure in snapper across the mid-west and south-west of WA.

### 4.3 Temporal variability in genetic structure

The analysis of the two temporally replicated samples uncovered differences in ancestry membership to the three identified stocks at Esperance, but not at Cockburn Sound. Temporal variation in population genetic structure has been reported for numerous marine species (e.g. Papetti et al. 2009, Hogan et al. 2010, Watts et al. 2010, Jackson et al. 2018, Quintero-Galvis et al. 2020). When comparing the repeated Esperance samples, there was a significant reduction in ancestry characteristic of our SA sample over time and an increased contribution from stocks further west (i.e., mid-west and south-west). This shift suggests that the genetic structure of snapper off Esperance may largely depend on dynamics of snapper populations in SA and along the south coast of WA, including population density and recruitment strength. Recent stock depletions and prolonged recruitment failure in the SA gulfs may account for the change in population genetic structure at Esperance (Fowler et al. 2020). Our results also suggest that the position of the genetic discontinuity detected in the south-east of our study range varies temporally. Overall, our temporal analyses indicate that incorporating samples collected at multiple timepoints in population genetic studies can reveal important biological information that may have otherwise gone undetected.

### 4.4 Suitability of current stock boundaries based on population genomics

Here, a population genomics framework has generated detailed information on the population dynamics of a highly mobile fishery important teleost. Employing a large dataset of genetic markers gave us sufficient power to uncover patterns of population genetic structure and connectivity at multiple spatial scales. We can therefore confidently use our findings to inform the spatial assessment and management of snapper. The patterns of genetic structure in snapper uncovered in this study partly contrast with current spatial scales of stock assessment and management. Across WA (outside of the inner gulfs of Shark Bay), three snapper stocks are currently recognized for assessment and management: the Shark Bay Oceanic (comprising the Gascoyne), West Coast (comprising Kalbarri, Lancelin, Cockburn Sound and Busselton) and South Coast stocks (comprising Albany and Esperance; see Figure 1). We detected a lack of genetic structure over an area extending ∼800 km between the Gascoyne and Lancelin. Our results therefore support a single management area across this region (i.e., the mid-west stock in this study).

In contrast to the mid-west, in the south-west of WA between Cockburn Sound and Albany (i.e., the south-west in this study), local-scale genetic structure was uncovered whereby genetic differentiation increased with coastal distance (i.e., an IBD pattern). Simulation work by Spies et al. (2015) showed that splitting an area characterised by IBD for management purposes leads to more favourable outcomes than if considered singularly.

Further, they demonstrated that the most optimal outcomes could be achieved by splitting a management region according to dispersal distance and fishing effort. Others suggest that under a pattern of IBD, spatial scales of management might be based on the x-intercept of spatial autocorrelation correlograms, a value that indicates the distance at which individuals become genetically independent (Diniz-Filho and De Campos Telles 2002, Östman et al. 2017). Considering our estimate of mean dispersal distance per generation determined from the IBD slope (∼400 km), the x-intercept of our spatial autocorrelation correlogram (∼300 km) and that fishing pressure is greater along the west coast near the Perth metropolitan region than the south coast (Gaughan and Santoro 2021), the current management boundary on the south-west corner of the state (at 115.5°E, see Figure 1) may be suitable.

Along the south coast of WA, the region between the south-west corner and the SA border is considered singularly for stock assessment and management purposes (i.e., the South Coast stock; Gaughan and Santoro 2021). Here we detected significant gene flow across this region. However, gene flow seems to vary temporally and therefore the boundary between SA and WA snapper populations is not stable and varies across space through time. Considering that the genetic structure of snapper off Esperance changes over time and that catch data indicate that the biomass of snapper in the south east of WA is likely low relative to adjacent areas (Norriss et al. 2016), it may be appropriate to consider locations in WA east of Albany separately for stock assessment and management purposes if the genetic composition of snapper in the Esperance region is not regularly reassessed. The south-coast presents a more complicated situation for spatial assessment and management and highlights the difficulties in translating complex biological processes into distinct units suitable for fisheries management. Population genomic work involving finer scale sampling along the south coast of WA and the west coast of SA (i.e., between Albany and Ceduna) would be valuable for gaining a more complete understanding of spatial variation in the contribution of SA genotypes to snapper along the south coast of WA and therefore for discerning the most suitable locations for drawing management boundaries.

## 5 CONCLUSIONS

The fluidity of the marine environment, along with the highly mobile and abundant nature of the species which occupy it, makes the characterisation of evolutionary and demographic patterns difficult. Our study highlights the utility of large datasets of genetic markers in improving understanding of spatial population structure and connectivity in marine species with high dispersal potential. The findings likely relate to variation in physical factors, such as local ocean circulation and coastal geomorphology, as well as in biological factors, such as timing of reproduction, spawning site preference and migratory behaviour. Overall, our study demonstrates the value of population genomics in assessing the suitability of the geographical scales at which stock assessment and management are carried out. Additionally, our work adds to the extensive body of literature showing that marine species are often characterised by complex population structure, rather than panmixia, and that population structure is not always static across time.

## DATA AVAILABILITY STATEMENT

The SNP dataset are available on figshare: doi to be provided upon acceptance.

## ACKNOWLEDGEMENTS

This work was financially supported by the Australian Research Council (LP180100756 to LBB), Flinders University and by the PhD scholarships to AB provided by AJ and IM Naylon and the Playford Trust. We sincerely thank those who assisted with the collection of tissue samples, which included staff from the Department of Primary Industries and Regional Development WA and the South Australian Research and Development Institute. We also thank Michelle Gardner for providing the temporal sample from Esperance. We thank Andrea Barceló and Diana-Elena Vornicu at the Molecular Ecology Lab Flinders University for their assistance in the laboratory.

---

**TABLE S1.** Biological and catch data for each of the 10 snapper samples (i.e., the nine Western Australian samples, including the two temporal samples, and the comparative sample from South Australia) as well as a summary of timing of spawning for each of the sites sampled. Values in the gonad stage column are means and standard deviations (in parentheses). Gonad stages are: 1, immature; 2, resting; 3, developing; 4, developed; 5, spawning; 6, spent.

|  | **Jurisdiction** | **Management area** | **Avg. latitude** | **Avg. longitude** | **Spawning season (peak)** | **Catch dates** | **Sector** | **Sex ratio (M:F)** | **Gonad stage** |
| --- | --- | --- | --- | --- | --- | --- | --- | --- | --- |
| **Gascoyne (GAS)** | Western Australia | Shark Bay Oceanic | -24.6667 | 113.0833 | May-Sep (Jun/Jul) | Jul 18 | Commercial | 0.6 | 4.0 (0.8) |
| **Kalbarri (KAL)** | Western Australia | West Coast | -27.8557 | 113.425 | Jul-Oct (Jul/Aug) | Jul-Sep 18 | Commercial/<br>Research | 1.2 | 3.6 (0.9) |
| **Lancelin (LAN)** | Western Australia | West Coast | -30.964 | 115.036 | Oct-Dec (Nov/Dec) | Oct 18 | Research | 1.1 | 3.3 (1.0) |
| **Cockburn Sound (CS)** | Western Australia | West Coast | -32.1855 | 115.7156 | Oct-Dec (Nov/Dec) | Oct 18 | Research | 1.2 | 4.4 (0.5) |
| **Cockburn Sound 14 (CS14)** | Western Australia | West Coast | -32.194 | 115.7442 | Oct-Dec (Nov/Dec) | Oct 14 | Research | 0.8 | NA |
| **Busselton (BUS)** | Western Australia | West Coast | -33.6343 | 115.2958 | Oct-Dec (Nov/Dec) | Jul-Aug 18 | Recreational | 1.0 | 2.5 (0.6) |
| **Albany (ALB)** | Western Australia | South Coast | -35.1815 | 118.3654 | Oct-Dec (Oct/Nov) | Aug-Nov 18 | Commercial | 0.9 | 3.5 (1.1) |
| **Esperance (ESP)** | Western Australia | South Coast | -34.2402 | 121.3276 | Oct-Dec (Oct/Nov) | Jun-Feb 19/20 | Commercial/<br>Recreational | NA | NA |
| **Esperance 10 (ESP10)** | Western Australia | South Coast | NA | NA | Oct-Dec (Oct/Nov) | Mar 10 | Recreational | NA | NA |
| **Ceduna (CED)** | South Australia | Spencer Gulf/<br>West Coast | -32.3 | 133.8 | Nov-Feb (Dec) | Sep 18, Aug 19 | Commercial | 1.1 | 1.6 (0.5) |

**TABLE S2.**
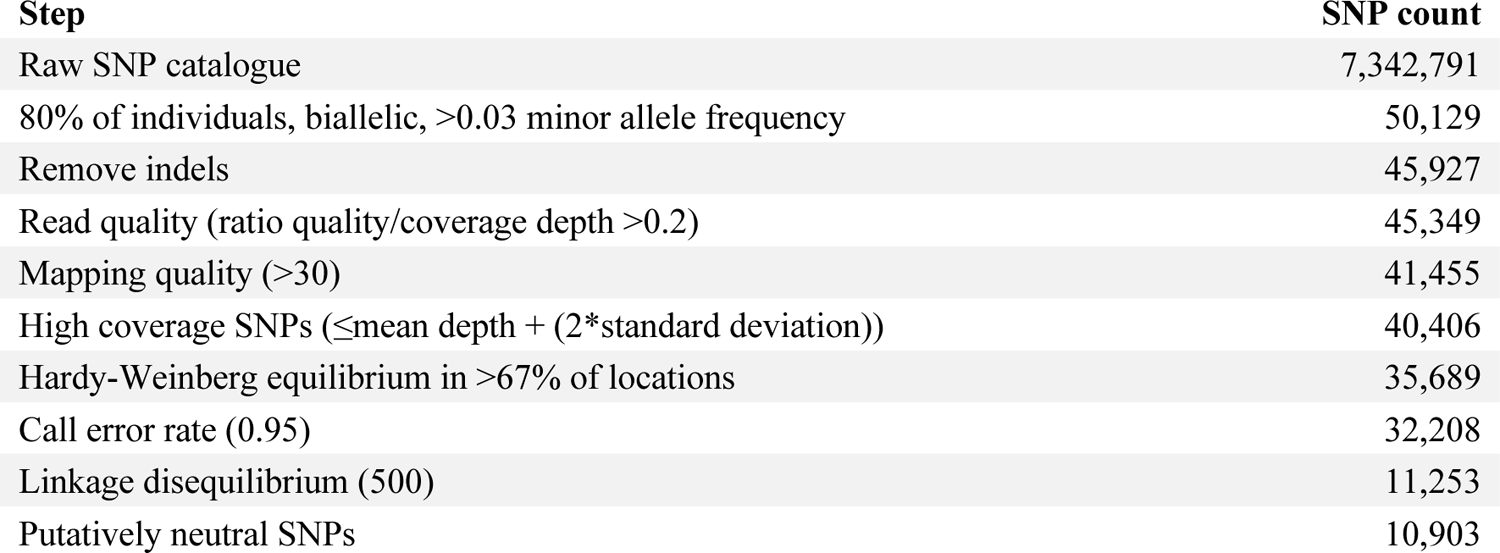
Numbers of SNPs retained after each bioinformatics filtering step.

| **Step** | **SNP count** |
| --- | --- |
| Raw SNP catalogue | 7,342,791 |
| 80% of individuals, biallelic, >0.03 minor allele frequency | 50,129 |
| Remove indels | 45,927 |
| Read quality (ratio quality/coverage depth >0.2) | 45,349 |
| Mapping quality (>30) | 41,455 |
| High coverage SNPs (≤mean depth + (2*standard deviation)) | 40,406 |
| Hardy-Weinberg equilibrium in >67% of locations | 35,689 |
| Call error rate (0.95) | 32,208 |
| Linkage disequilibrium (500) | 11,253 |
| Putatively neutral SNPs | 10,903 |

